# HIV envelope trimers and gp120 as immunogens to induce broadly neutralizing antibodies in cows

**DOI:** 10.1101/2024.03.20.585065

**Authors:** Pilar X. Altman, Mara Parren, Huldah Sang, Gabriel Ozorowski, Wen-Hsin Lee, Vaughn V. Smider, Ian A. Wilson, Andrew B. Ward, Waithaka Mwangi, Dennis R. Burton, Devin Sok

**Affiliations:** Department of Immunology and Microbiology, The Scripps Research Institute, La Jolla, CA 92037, USA; Consortium for HIV/AIDS Vaccine Development (CHAVD), The Scripps Research Institute, La Jolla, CA 92037, USA; IAVI Neutralizing Antibody Center, The Scripps Research Institute, La Jolla, CA 92037, USA; Department of Diagnostic Medicine/Pathobiology, College of Veterinary Medical, Kansas State University, Manhattan, Kansas 66506, USA; Department of Integrative Structural and Computational Biology, The Scripps Research Institute, La Jolla, CA 92037, USA; Department of Molecular Medicine, The Scripps Research Institute, La Jolla, CA, 92037, USA; Applied Biomedical Science Institute, San Diego, CA, 92127, USA; Skaggs Institute for Chemical Biology, The Scripps Research Institute, La Jolla, CA, 92037, USA; Ragon Institute of Massachusetts General Hospital, Massachusetts Institute of Technology, and Harvard University, Cambridge, MA 02139, USA; International AIDS Vaccine Initiative, New York, NY 10004, USA

## Abstract

The study of immunogens capable of eliciting broadly neutralizing antibodies (bnAbs) is crucial for the development of an HIV vaccine. To date, only cows, making use of their ultralong CDRH3 loops, have reliably elicited bnAbs following immunization with HIV Envelope trimers. Antibody responses to the CD4 binding site have been readily elicited by immunization of cows with a stabilized Env trimer of the BG505 strain and, with more difficulty, to the V2-apex region of Env with a cocktail of trimers. Here, we sought to determine whether the BG505 Env trimer could be engineered to generate new bnAb specificities in cows. Since the cow CD4 binding site bnAbs bind to monomeric BG505 gp120, we also sought to determine whether gp120 immunization alone might be sufficient to induce bnAbs. We found that engineering the CD4 binding site by mutation of a key binding residue of BG505 HIV Env resulted in a reduced bnAb response that took more immunizations to develop. Monoclonal antibodies isolated from one animal were directed to the V2-apex, suggesting a re-focusing of the bnAb response. Immunization with monomeric BG505 g120 generated no serum bnAb responses, indicating that the ultralong CDRH3 bnAbs are only elicited in the context of the trimer in the absence of many other less restrictive epitopes presented on monomeric gp120. The results support the notion of a hierarchy of epitopes on HIV Env and suggest that, even with the presence in the cow repertoire of ultralong CDRH3s, bnAb epitopes are relatively disfavored.

## Introduction

The enormous diversity of HIV suggests that a vaccine should elicit broadly neutralizing antibodies (bnAbs) capable of neutralizing a wide array of HIV viral strains [1–5]. Many bnAbs have been isolated from HIV-infected donors, and they have helped define vaccine targets on HIV Envelope (Env) [6, 7]. However, eliciting bnAbs is difficult due to several characteristic features, including high levels of somatic hypermutation, specific V_H_ gene usage, insertions, deletions, and long heavy chain complementary determining (CDRH3) region [8–10]. The latter feature is relatively rare in the naïve repertoires of humans and virtually all animal models, with the exception of cows, creating difficulty in initiating bnAb responses. The CDRH3s of the cow antibody repertoire are, on average, 26 amino acids in length, as compared to the average 15 amino acids in the human repertoire [11–21]. In addition, they have a small subset (about 10% of the repertoire) of ultralong CDRH3s [16, 21–24]. For the purpose of this study, we categorize antibodies based on their CDRH3 length as follows: “ultralong” antibodies as having CDRH3 lengths ≥50 amino acids, “long” cow antibodies with CDRH3 lengths ranging from 25-49 amino acids, and “short” cow antibodies as those with CDRH3 lengths <25 amino acids.

Early studies immunized pregnant cows with non-well-ordered HIV Env trimers, yielding cross-clade HIV neutralizing colostrum, which predominately targeted the CD4 binding site (CD4bs) [25, 26]. Later studies showed that cows could elicit broadly neutralizing antibodies reliably and rapidly to both the CD4bs and V2-apex when immunized with well-ordered trimer immunogens [27–29]. One of the CD4bs studies showed that, in cows, a single immunogen was able to rapidly elicit a potent and broad broadly neutralizing antibody response directed towards the CD4bs [27]. Initially, it was unclear if broadly neutralizing antibody responses could develop to target epitopes beyond the CD4bs, given that the response to this site was so overwhelming. However, we found that using one or multiple V2-apex bnAb-focusing trimer immunogens as well as a cocktail of trimers, we could elicit a cross-clade and somewhat potent bnAb response that targeted the V2-apex in two cohorts of cows [28]. Some questions remained unanswered. First, could we re-direct the CD4bs cow bnAb by simply making it less accessible to antibody recognition by mutation. Second, was a trimer immunogen absolutely required to elicit bnAbs in cows since the cow bnAbs did bind to monomeric gp120. In other words, did one need to provide a protein in which the only way to access the bnAb site, in this case the CD4bs, was through deployment of an ultralong CDRH3 or would the presentation of the site in a less restricted context on gp120 nevertheless induce a response capable of recognizing the trimer.

To investigate the first question, we immunized cows with a modified version of the immunogen, BG505 D368R SOSIP.664 trimer, (SOSIP.664 will be referred to as SOSIP from this point forward). This epitope-mutated SOSIP was designed to knock out a residue important for binding and neutralization by CD4bs bnAbs [30, 31]. To investigate the second question, we immunized cows with BG505 gp120 that has the same sequence in the gp120 region as BG505 SOSIP trimer. Following repeated immunization, serum neutralization was evaluated longitudinally, and IgG+ B cells were collected and sorted from cows with relevant neutralizing responses. From the isolated cells, we identified and characterized bnAbs, especially in comparison to other known cow bnAbs.

## Results

### Cows immunized with BG505 D368R SOSIP had a late and reduced cross-clade neutralizing response compared to those immunized with wild type BG505 SOSIP, while cows immunized with BG505 gp120 had no cross-clade neutralizing response

In this study, three groups of two cows were immunized with either BG505 SOSIP, BG505 D368R SOSIP, or BG505 gp120. Group 1, comprising cow-2651 and cow-2518, was immunized with BG505 SOSIP, and Group 2, consisting of cow-2594 and cow-2604, was immunized with BG505 D368R SOSIP. Both groups were immunized on day 0, followed by five boosts on days 36, 76, 118, 258, and 384 for a total of six immunizations. Group 3 cows, cow-596 and cow-2541, were immunized with BG505 gp120 on day 0 and boosted four times on days 36, 76, 118, and 258, culminating in five immunization events. Each immunization used 200 µg of immunogen per animal. Serum samples and PBMCs were collected for analysis over the course of the immunizations. The immunization protocols are outlined in Fig 1.

**Fig. 1.**
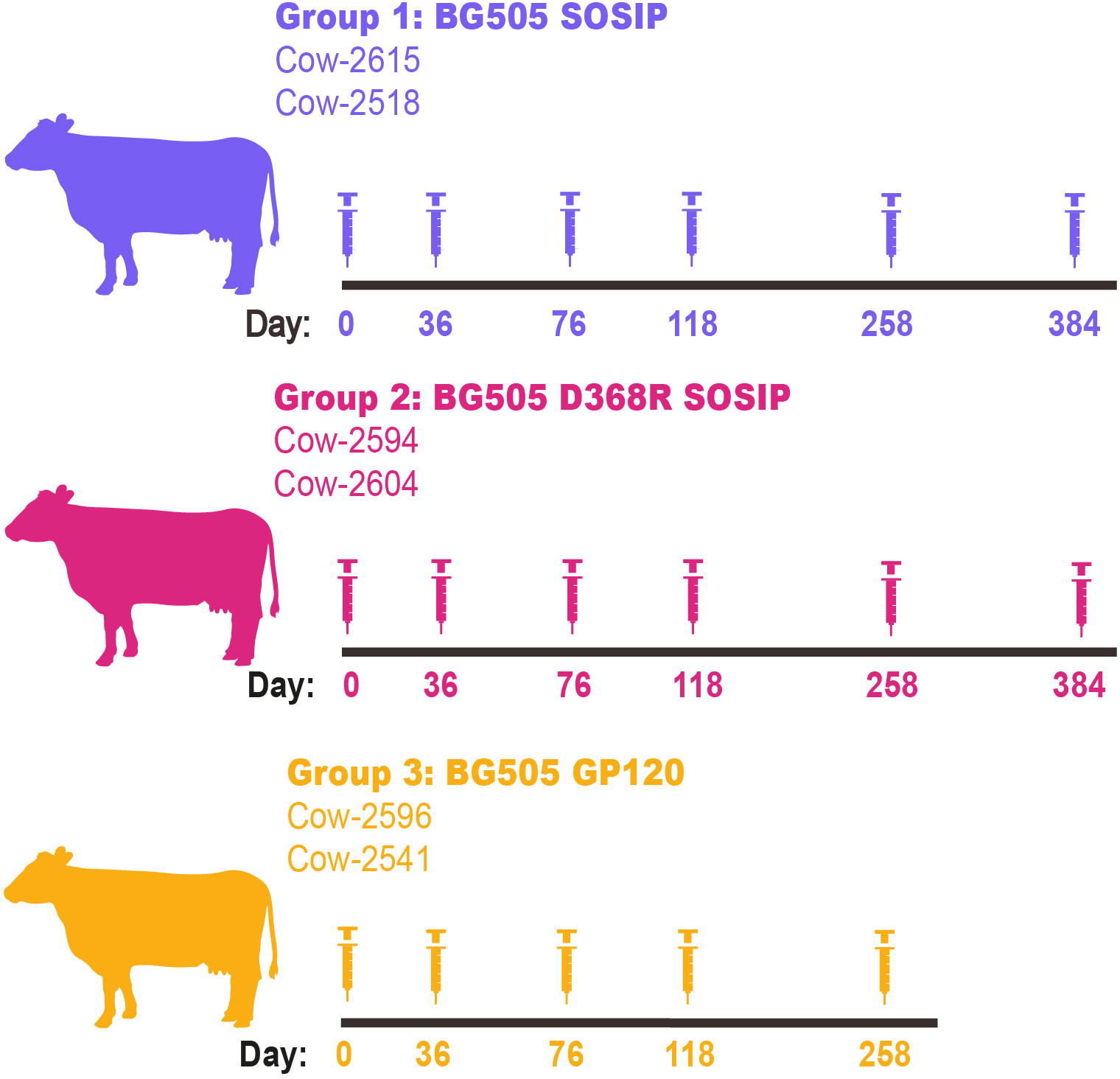
Immunization of cows with BG505 recombinant proteins. Three groups of cows are grouped by the recombinant protein they were immunized with. Group 1 (top) immunized with BGS0S SOSIP is shown in purple. Group 2 (middle) was immunized with BGS0S D368R SOSIP and is shown in pink. Group 3 (bottom) immunized with BGS0S gp120 is shown in yellow

Serum samples from 12 time points across the immunization schedule were purified to isolate IgG and minimize non-specific background activity. Initial serum testing was evaluated for the presence and potency of a neutralizing response against the autologous wild-type BG505 pseudovirus (Table 1). In Group 1, both cows developed a similar neutralizing response of comparable potency and timing; cow-2518 developed a detectable response by day 85, and cow-2615 developed a detectable response by day 91. The neutralizing responses were enhanced over the course of successive immunizations and reached a maximum by the final time point. Group 2 cows also developed neutralization to BG505 pseudovirus; cow-2604 was able to weakly neutralize BG505 in as little as 85 days, but the response did not strengthen until after the fifth immunization. Cow-2594 did not display an autologous neutralizing response until day 265. By the final time point, it matched the potency observed in the BG505 SOSIP immunized cows. Cows from Group 3 did not develop a detectable neutralizing response to BG505 over the entire period of monitoring.

**Table 1.** Neutralization of BG505 isolate for all cows in this study. Neutralization ID_50_ titers are shown for IgG purified from sera from all cows against the BG505 isolate over the course of the study. ID_50_ values are shown as 1/dilution. Values marked with X are not available.

| ID<sub>50</sub><br>(1/dilution) | Group 1<br>BG505 SOSIP |  | Group 2<br>BG505 D368R SOSIP |  | Group 3<br>BG505 gp120 |  |
| --- | --- | --- | --- | --- | --- | --- |
| Cow | 2615 | 2518 | 2594 | 2604 | 2596 | 2541 |
| Day 0 | <35 | <35 | <35 | <35 | <35 | <35 |
| Day 13 | <35 | <35 | <35 | <35 | <35 | <35 |
| Day 46 | <35 | <35 | <35 | <35 | <35 | <35 |
| Day 49 | <35 | <35 | <35 | <35 | <35 | <35 |
| Day 85 | <35 | 263 | <35 | 78 | <35 | <35 |
| Day 91 | 66 | 331 | <35 | 53 | <35 | <35 |
| Day 127 | 434 | 982 | <35 | 87 | <35 | <35 |
| Day 133 | 442 | 1194 | <35 | 106 | <35 | <35 |
| Day 265 | 3019 | 1161 | 806 | <35 | <35 | <35 |
| Day 273 | 6753 | 2403 | 860 | 211 | <35 | <35 |
| Day 384 | 3388 | 2047 | 489 | 2277 | X | X |
| Day 398 | 6892 | 11225 | 11687 | 3670 | X | X |

To evaluate the breadth of neutralizing activity at the final time point, sera were tested against a 12-virus global panel [32] (Table 2). Group 1 cows elicited a neutralizing response to 92% and 100% of the panel viruses by day 384. Group 2 exhibited a less broad response and only neutralized 50% and 67% of the viruses in the panel by day 384. Group 3 had one cow neutralize one virus (8% breadth), specifically the ultra-sensitive virus 398F1 from clade A, on day 273.

**Table 2.** Neutralization titers for all cows in this study at the terminal timepoint. Neutralization ID_50_ titers are shown for IgG purified from sera from all cows at the terminal timepoint tested against the 12-virus global panel. Geomean ID_50_ and percent breadth are shown for the 12-virus global panel at the bottom. ID_50_ values are shown as 1/dilution.

| ID<sub>50</sub> (1/dilution) |  | Group 1<br>BG505 SOSIP |  | Group 2<br>BG505 D368R SOSIP |  | Group 3<br>BG505 gp120 |  |
| --- | --- | --- | --- | --- | --- | --- | --- |
| Cow |  | 2615 | 2518 | 2594 | 2604 | 2596 | 2541 |
| Virus | Clade | Day 398 |  | Day 398 |  | Day 273 |  |
| MLV | N/A | <35 | <35 | <35 | <35 | <35 | <35 |
| 246F3 | AC | 109 | 794 | 3757 | 163 | <35 | <35 |
| 25710 | C | 2178 | 419 | 183 | 1254 | <35 | <35 |
| 398F1 | A | 4504 | 9425 | 8963 | 122 | 261 | <35 |
| BJOX2000 | BC | 455 | 620 | 341 | 530 | <35 | <35 |
| CE1176 | C | 436 | 2566 | <35 | <35 | <35 | <35 |
| CEO217 | C | 2305 | 4398 | <35 | <35 | <35 | <35 |
| CH119 | BC | <35 | 289 | <35 | <35 | <35 | <35 |
| CNE55 | AE | 331 | 633 | <35 | 267 | <35 | <35 |
| CNE8 | AE | 211 | 1117 | <35 | 363 | <35 | <35 |
| TRO.11 | B | 74 | 87 | 620 | 164 | <35 | <35 |
| X1632 | G | 2031 | 2953 | 11508 | 84 | <35 | <35 |
| X2278 | B | 452 | 599 | <35 | <35 | <35 | <35 |
| Geomean ID<sub>50</sub> |  | 576 | 969 | 1571 | 255 | 261 | <35 |
| % Breadth |  | 92% | 100% | 50% | 67% | 8% | 0% |

Following the confirmation of cross-clade neutralization, we next wanted to determine when these titers developed in relation to the immunization time points. Using time points about two weeks after each immunization as well as a pre-bleed, we tested the full 12-virus panel and MLV for neutralization (Fig 2). Group 1 animals developed consistent weak heterologous neutralizing activity by day 85 (8% breadth, n=12). The breadth of neutralization of the response for these cows developed more slowly than in the previous study (Sok *et al.* 2017) using a somewhat different immunization schedule and adjuvant administration [27]. Group 2 animals developed weak heterologous neutralization at day 85 to 398F1, but breadth did not increase until after the fifth immunization on day 258 and potency was only slightly strengthened after the sixth immunization. Group 3 cows exhibited weak neutralization of virus 398F1, with one of the cows losing the response by the final time point.

**Fig 2.**
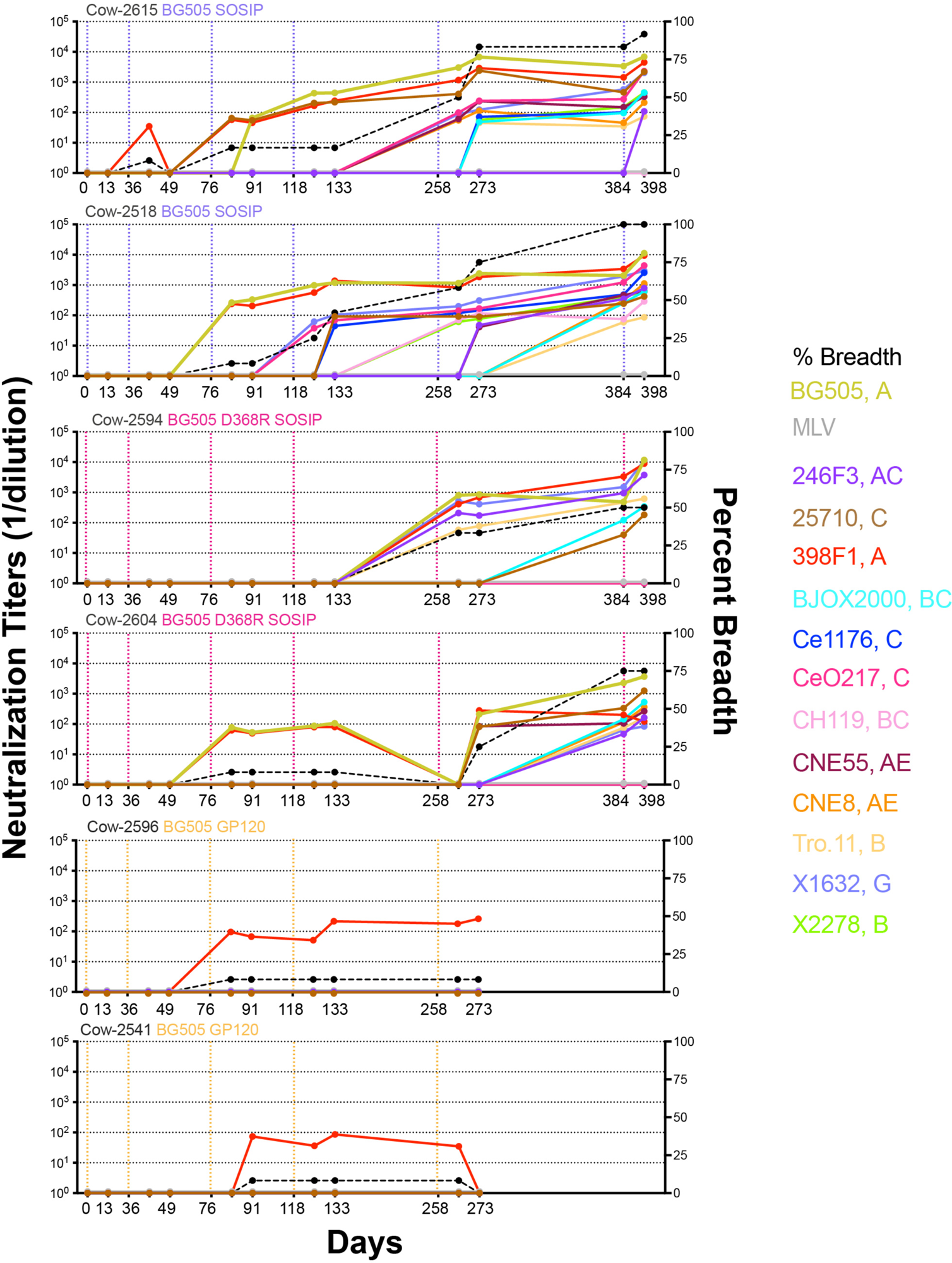
Neutralization of a 12-virus panel by sera from different time points following immunization shows cross-clade neutralizing responses with recombinant trimers but not monomers. Longitudinal samples were collected and tested for all six cows on a 12-virus global panel as well as on BG505 and MLV. Neutralization ID_50_ titers are shown by dots and each virus is labeled a different color. Vertical dotted lines represent immunization prime and boost dates. BG505 SOSIP is shown in purple, BG505 D368R SOSIP in pink, and BG505 gp120 in yellow. Neutralization percent breadth is represented by a dashed black line whose y-axis is shown on the right and only includes the breadth of the 12-virus global panel. ID_50_ values are shown as 1/dilution on the y-axis. Neutralization was only recorded for those samples that neutralized at dilutions greater than 35-fold.

We next sought to define the specificities of serum antibodies from the immunized animals by investigation of competition with bnAbs NC-Cow1, VRC01, PGT121, 35O22, PGDM1400, and CAP256v9 (Table 3, Fig 3). We found Group 1 cows to have varying degrees of competition with different bnAbs. Cow-2615 had the most marked competition with bnAb 35022 at the gp120/gp41 interface and some competition with the other bnAbs, most notably CD4bs and V2-apex bnAbs. Cow-2518 had marked competition (<70%) with all the bnAbs tested by the final time point. Group 2 cows displayed strongest competition with the gp120/gp41 interface bnAb and some competition with the other bnAbs. For all cows, the degree of competition increased over the course of immunization. For cow-2594 and cow-2604, 35O22 competition started at low levels in as little as 91 days, but further competition at other epitopes expanded after the final immunization.

**Fig 3.**
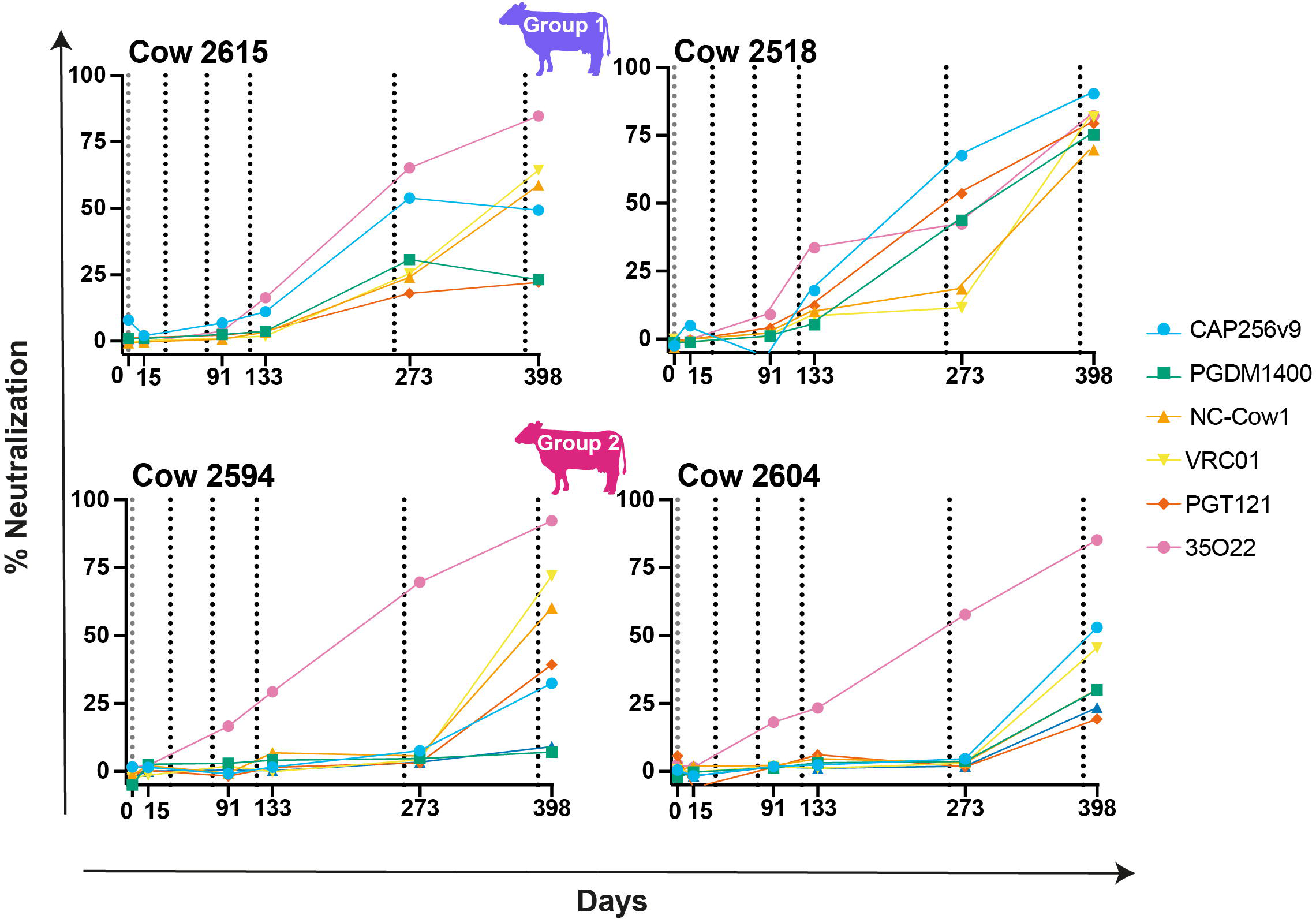
BG505 WT SOSIP binding competition in four SOSIP immunized cows followed over time. Longitudinal samples were collected and tested for Group 1 and Group 2 cows to test competition binding with known antibodies. Each dot represents percent neutralization at that time point with the antibody of the corresponding color. Vertical dotted lines represent immunizations.

**Table 3.** Mapping sera epitope specificity at the terminal timepoint. Neutralization ID_50_ titers are shown for IgG purified from sera from all cows in the study at the terminal timepoint tested against the 12-virus global panel. Geomean ID_50_ and percent breadth are shown for the 12-virus global panel at the bottom. ID_50_ values are shown as 1/dilution.

| Percent Competition |  | Group 1<br>BG505 SOSIP |  | Group 2<br>BG505 D368R SOSIP |  |
| --- | --- | --- | --- | --- | --- |
| Cow |  | 2615 | 2518 | 2594 | 2604 |
|  |  | Day 398 |  |  |  |
| CD4bs | NC-Cow1 | 59% | 70% | 60% | 30% |
|  | VRC01 | 64% | 82% | 72% | 46% |
| V3 glycan | PGT121 | 22% | 79% | 39% | 19% |
| gp120/gp41 interface | 35O22 | 85% | 82% | 92% | 85% |
| V2-Apex | PGDM1400 | 23% | 75% | 7% | 30% |
|  | CAP256v9 | 49% | 90% | 32% | 53% |

### Isolation of monoclonal antibodies from cow-2594 and cow-2604

After confirming that neutralization within the SOSIP immunized groups, we shifted our efforts to isolating monoclonal antibodies. We chose to focus on cow-2594 and cow-2604, as BG505 SOSIP-immunized cows have been studied extensively in Sok *et al*. 2017, and BG505 gp120-immunized cows had negligible cross-clade neutralizing responses [27]. We used two rounds of single IgG^+^ B Cell sorting using an epitope specific sorting strategy. The sorts used PBMCs from the final time point day 398 stained with goat anti-cow IgG conjugated with FITC and biotinylated antigens conjugated to streptavidin fluorophores to isolate the B-cell populations of interest (Fig 4). All SOSIP baits were biotinylated and stained with streptavidin conjugated to fluorophores. For Sort 1, we used BG505 WT SOSIP and BG505 SOSIP complexed with four known Fabs targeting different epitopes, including PG04 (CD4bs), PGT145 (V2-Apex), PGT151 (MPER), and PGT121 (V3-glyan). Both BG505 baits were coupled to separate fluorophores to distinguish those that were only BG505 (PE) and those that were complexed with Fab (AF 647). B cells were sorted if they were bound to BG505 WT but not the BG505-Fab complex, indicating that they were competing with the bound Fab. Sort 2 was done using only final cow-2604 samples, utilizing BG505 and 25710 SOSIP as baits. Cells were sorted based on whether they were bound to both BG505 SOSIP coupled with BV421 and BG505 coupled with BUV737 and bound to 25710 SOSIP coupled with PE. Single-cell PCR was used to isolate and amplify the variable regions from sorted B cells, which were later cloned into human heavy chain and lambda light chain antibody expression vectors and sequenced as previously described [27]. During the first sort, the majority of isolated cells from cow-2604 were PGT121 competing, while cow-2594 had predominantly cells sorted from the PGV04 complexed bait (Table 4). Despite a similar number of cells sorted, cow-2604 had a higher rate of heavy chain recovery. A majority of these recovered CDRH3s from cow-2604 were ultralong, while cow-2594 had mostly short CDRH3s recovered. Sort 2 had a slight predominance of short CDRH3s recovered for cow-2604. Our recovery results suggest an enrichment of ultralong CDRH3s in the trimer-immunogen binding pools as compared to the total cow repertoire, where the ultralong antibodies account for about 10% of total CDRH3s [16, 21–24].

**Fig 4.**
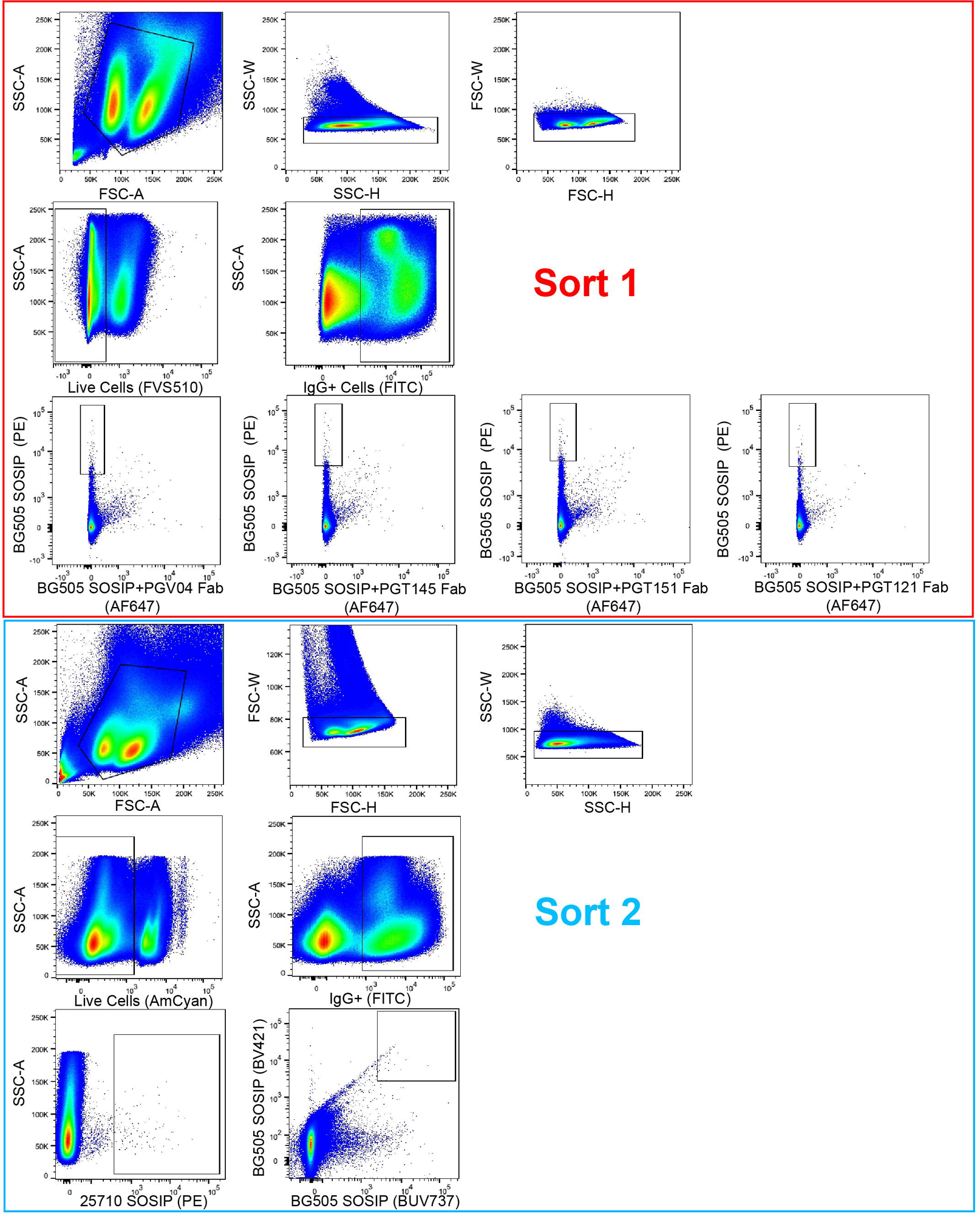
Graphs of two single B cell sorts used to isolate monoclonal antibodies of interest. Sort 1 was done on terminal time point samples from BGS0S D368R SOSIP-immunized cows, cow-2594 and cow-2604 (top). Sort 2 was only done on cow-2604 samples (bottom). Approximate gating is shown for each graph. Fluorophores are shown on the appropriate axes in parenthesis.

**Table 4.** Summary of results from Sort 1 and Sort 2. Total cells and recovered heavy chains are shown for each cow. Cells for Sort1 are sorted by Fab-BG505 complex and recovered heavy chains are sorted by CDRH3 length category. NA stands for not available as those experiments were not performed for that sample.

| Summary of Sort Results |  | Sort 1 |  |  |  |  | Sort 2 |
| --- | --- | --- | --- | --- | --- | --- | --- |
|  |  | Total | PGT151 | PGV04 | PGT121 | PGT145 | Total |
| Cow-2594 | Total Cells Sorted | 390 | 95 | 145 | 63 | 87 | NA |
|  | PCR Wells Positive for Heavy Chains | 24 | 3 | 8 | 8 | 5 | NA |
|  | Heavy Chain Sequences Recovered and Tested | 23 | 3 | 7 | 8 | 5 | NA |
|  | Short CDRH3 (0-24) | 14 | 2 | 2 | 6 | 4 | NA |
|  | Long CDRH3 (25-49) | 5 | 0 | 4 | 1 | 0 | NA |
|  | Ultralong CDRH3 (49+) | 3 | 1 | 0 | 1 | 1 | NA |
| Cow-2604 | Total Cells Sorted | 400 | 64 | 78 | 176 | 82 | 462 |
|  | PCR Wells Positive for Heavy Chains | 91 | 20 | 32 | 19 | 20 | 40 |
|  | Heavy Chain Sequences Recovered and Tested | 78 | 18 | 27 | 17 | 16 | 36 |
|  | Short CDRH3 (0-24) | 27 | 3 | 10 | 11 | 3 | 20 |
|  | Long CDRH3 (25-49) | 5 | 1 | 1 | 3 | 0 | 1 |
|  | Ultralong CDRH3 (49+) | 46 | 14 | 16 | 3 | 13 | 15 |

After the isolation process, we recombinantly expressed the isolated monoclonal antibodies. Heavy chains were expressed with their native light chain if it was recovered, and/or V30, a universal cow light chain that predominately pairs with ultralong CDRH3s [15]. The V30 pairing was specifically applied to heavy chains derived from the IGHV1-7 gene to ensure as thorough testing as possible. Long CDRH3s derived from the IGHV-1 gene are not always able to pair with the universal light chain but were tested without regard to length. Antibodies were screened using a high-throughput Expi293 expression system, with antibody expression and binding to BG505 SOSIP conducted on unpurified supernatant using ELISA assays. Antibodies positive for expression were evaluated for neutralization of BG505, 25710, X1632, and 246F3 pseudoviruses (Table 5). Briefly, cow-2594 had a total of 20 heavy chains paired with light chains and four heavy chains paired with the universal light chain. 12 of the heavy chains had CDRH3s that were short, five were long, and three were ultralong for cow-2594 tested with their native light chain. Of those paired with a universal light chain, none were short, one was long, and three were ultralong. For cow-2604, a total of 95 monoclonal antibodies (mAbs) were tested with their native light chain, and 63 mAbs were assessed with the universal light chain. In the native light chain set, 37 of the heavy chain CDRH3s were short, six were long, and 52 were ultralong. Of those tested with the universal light chain, there were two long antibodies, 61 ultralong antibodies, and no short antibodies. Despite testing over a hundred different monoclonal antibodies, only two antibodies with cross-clade neutralization were isolated, both from cow-2604, Moo1 and Moo2 (Fig 5A).

**Fig 5.**
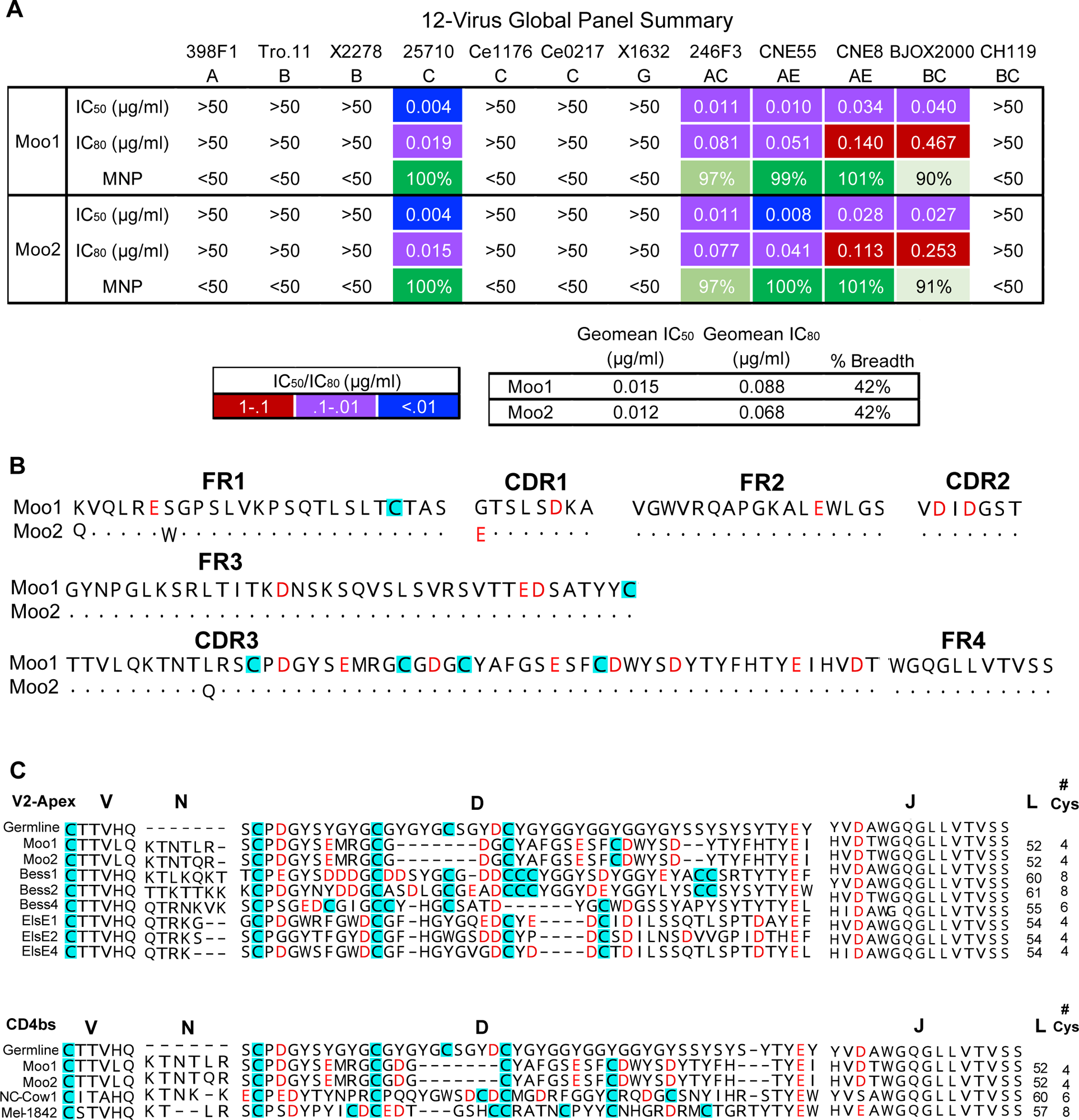
Broadly neutralizing antibodies Moo1 and Moo2 have ultralong CDRH3s and medium neutralization breadth and high potency. **(A)** Results of Moo1 and Moo2 mAbs measured against a 12-virus global panel for neutralization. Neutralizing IC50 and IC80 titers are shown for viruses neutralized at IC50 <50 μg/ml. Neutralizing geomean IC50 and geomean IC80 titers are shown for viruses neutralized at IC50 <50 μg/ml. Percent breadth on the panel is shown for each antibody, as well as the geomean maximum percent neutralization (MNP) for viruses neutralized at IC50 <50 μg/ml. **(B)** The variable regions of Moo1 and Moo2 are shown. Dots represent an identical amino acid at a given position. Negatively charged D and E amino acids are colored in red. Cysteines are highlighted in blue. **(C)** Moo1 and Moo2 amino acid CDRH3s are aligned with V2-apex cow bnAbs (top) and CD4bs bnAbs (bottom) as well as the germline VH1–7, DH8-2, and JH10 regions. CDRH3 lengths (L) and the number of cysteines in the D region of the CDRH3 are shown to the left of each sequence. Color coding is labeled in part A.

**Table 5.** Summary of screening results from Sort 1 and 2. Total cells and recovered heavy chains for each cow are shown. Cells for Sort1 are sorted by Fab-BG505 complex and recovered heavy chains are sorted by CDRH3 length category. NA stands for not available as those experiments were not performed for that sample.

| Summary of Screening |  |  | Sort 1 | Sort 2 | Short CDRH3<br>(0-24 AA) | Long CDRH3<br>(25-49 AA) | Ultralong CDRH3<br>(50+ AA) |
| --- | --- | --- | --- | --- | --- | --- | --- |
| Cow-2594 | All | Cells Sorted | 390 | NA | NA | NA | NA |
|  |  | PCR Wells Positive for Heavy Chains | 24 | NA | NA | NA | NA |
|  |  | Heavy Chain Sequences Recovered | 23 | NA | 14 | 6 | 3 |
|  | Universal Light Chain | IGHV-1*7 Heavy Chains | 4 | NA | 0 | 1 | 3 |
|  |  | Expressed in Screen | 0 | NA | 0 | 0 | 0 |
|  |  | Positive for BG505 SOSIP Binding | 0 | NA | 0 | 0 | 0 |
|  |  | mAb with Cross-Clade Neutralization | 0 | NA | 0 | 0 | 0 |
|  | Native Light Chains | Heavy Chain/Light Chain Pairs | 20 | NA | 12 | 5 | 3 |
|  |  | Expressed in Screen | 17 | NA | 10 | 5 | 2 |
|  |  | Positive for BG505 SOSIP Binding | 8 | NA | 5 | 2 | 1 |
|  |  | mAbs with Cross-Clade Neutralization | 0 | NA | 0 | 0 | 0 |
| Cow-2604 | All | Cells Sorted | 400 | 462 | NA | NA | NA |
|  |  | PCR Wells Positive for Heavy Chains | 91 | 40 | NA | NA | NA |
|  |  | Heavy Chain Sequences Recovered | 78 | 36 | 47 | 6 | 61 |
|  | Universal Light Chain | IGHV-1*7 Heavy Chains | 48 | 15 | 0 | 2 | 61 |
|  |  | Expressed in Screen | 44 | 13 | 0 | 2 | 55 |
|  |  | Positive for BG505 SOSIP Binding | 34 | 11 | 0 | 0 | 45 |
|  |  | mAbs with Cross-Clade Neutralization | 1 | 1 | 0 | 0 | 2 |
|  | Native Light Chain | Heavy Chain/Light Chain Pairs | 66 | 29 | 37 | 6 | 52 |
|  |  | Expressed in Screen | 56 | 25 | 32 | 5 | 44 |
|  |  | Positive for BG505 SOSIP Binding | 40 | 13 | 17 | 1 | 35 |
|  |  | mAbs with Cross-Clade Neutralization | 1 | 0 | 0 | 0 | 1 |

Moo1 was evaluated initially with both native light chain and universal light chain, but due to similar potency, the universal light chain was utilized for the rest of the experiments. We used the 12-virus global panel to test the neutralization potency, and unsurprisingly, Moo1 and Moo2 neutralized with similar potency and maximum neutralization percentage (Fig 5A). The Abs neutralized the same viruses, with Moo1 exhibiting a geomean IC_50_ of 0.015 µg/ml and Moo2 a geomean IC_50_ of 0.011 µg/ml with 42% breadth. The neutralization pattern was similar to that of Bess1 and Bess8 [28], though more potent than Bess8 and lacking neutralization of isolate X1632 >50 µg/ml. They all also displayed incomplete neutralization (<95%) of isolate BJOX2000, which is not uncommon among trimer-specific antibodies and was seen in Bess and ElsE bnAbs [28, 33–37]. Overall, the Moo antibodies were moderately broad and highly potent but less so than the best of the Bess, ElsE, NC-Cow, and Mel antibodies [27–29].

### Broadly neutralizing antibodies from cow-2604 contain ultralong CDRH3s

The two isolated bnAbs, Moo1 and Moo2, both contain ultralong CDRH3s.

They have minimal differences in their variable regions, with only one amino acid difference in their CDRH3, two differences in the FR1, and one difference in the CDR1 (Fig 5B). We looked at the amino acid sequence comparisons of Moo1 and Moo2 CDRH3 with the ultralong germline sequence and representative HIV cow bnAbs ElsE and Bess to the V2-apex from our earlier study, Altman et al. 2024, and also a comparison to germline and CD4bs cow bnAbs from Sok *et al.* 2017, and Heydarchi *et al.* 2022 [27–29] (Fig 5C). Moo1 and Moo2 had the same number of cysteines as ElsE antibodies and did retain three of the germline cysteines that are conserved in the germline across all the bnAbs. In the area of the “knob” region of CDRH3 and, likely, the area that interacts with antigen, there is not much similarity in amino acids, even among NC-Cow1, which came from a BG505 SOSIP-immunized cow. Comparison of Moo1 and Moo2 sequences show limited similarity with other known cow bnAbs at the amino acid level of the CDRH3.

### Epitope specificity of V2-apex targeting Moo antibodies and functional characterization

We used BLI competition assays with a panel of human bnAbs binding to BG505 WT SOSIP to determine Moo1 and Moo2 specificities (Fig 6A). Significant competition was observed with bnAb CAP256-VRC26.9 (87-95%) and with bnAb PGT145 (68-75%), indicating specificity for the V2-apex bnAb site. We further confirmed the V2-apex binding site by negative stain EM imaging of Moo1 Fab bound to BG505, along with RM20A3 base binding Fab added to increase angular sampling (Fig 6B).

**Fig 6.**
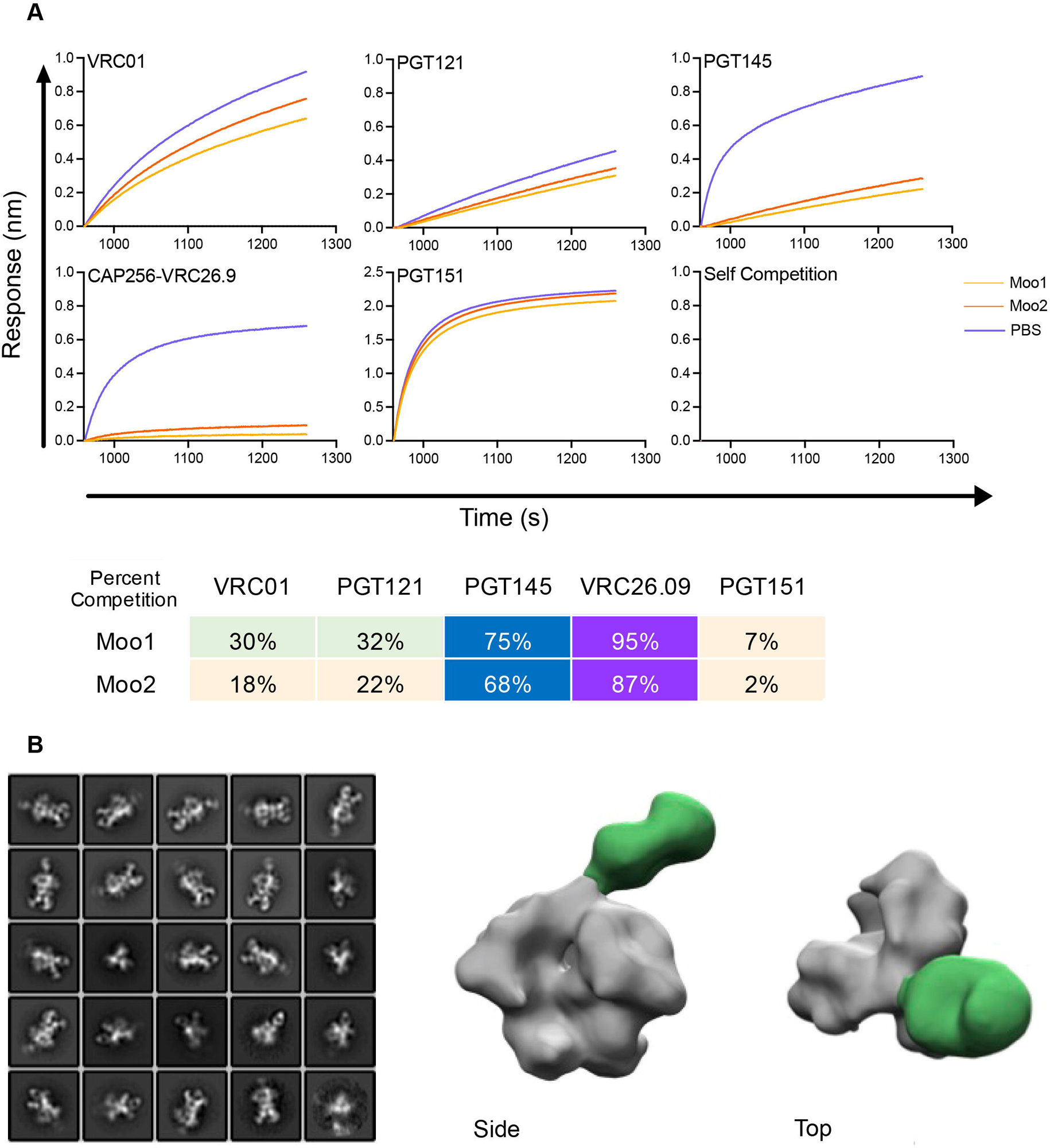
Broadly neutralizing antibodies Moo1 and Moo2 target the V2-apex epitope on the surface of HIV Env. **(A)** Moo1 and Moo2 were mapped on isolate BG505 SOSIP using BLI epitope binning. Antibodies VRC01 (CD4bs), PGT121 (V3 glycan), PGT145 (V2-Apex), CAP256-VRC26.9 (V2-apex), and PGT151 (MPER) as well as self-competition were used to represent the various HIV epitopes. The graphs (top) and calculated percent competition (bottom) are shown. There was evidence of competition at the V2-apex **(B)** Negative stain EM 2D classes (left) and 3D reconstruction (right) of Moo1 in complex with BGS0S SOSIP and base-binding Fab RM20A3. The 3D map is segmented with Moo1 colored green and RM20A3 (used to increase angular sampling) hidden for clarity.

To compare the characteristics of Moo antibodies with other human and cow V2-apex targeting bnAbs, we evaluated shared characteristics. First, we tested the Moo1 neutralization in the presence of V2-apex epitope mutations and noted that potency was significantly weakened by substitutions at positions N156, N160, and R166, which are residues important for the neutralization of HIV at the V2-apex for human and cow bnAbs (Fig 7A). Second, noting that many V2-apex bnAbs are trimer-specific or trimer-preferring, we compared Moo1 binding to BG505 SOSIP and gp120; no binding by ELISA was found for gp120 consistent with trimer specificity (Fig 7B). Third, we investigated the importance of tyrosine sulfation for Moo antibody binding to SOSIP trimer since this modification is known to have a significant role in the binding of many human V2-apex bnAbs, although the role of sulfation in the cow ElsE and Bess lineage antibodies is variable [38]. We used BLI and human AHC sensors to immobilize Moo1 and Moo2 and control antibodies. Sensors with immobilized antibodies were then exposed to mouse anti-sulfotyrosine antibodies to observe binding and evaluate tyrosine sulfation (Fig 7C). Moo1 and Moo2 did not bind to mouse anti-sulfotyrosine antibodies despite DY and DXY motifs in the CDRH3 (Fig 5B and C). In summary, Moo antibodies bind to the V2-apex of HIV and are unable to bind to recombinant monomer protein similar to many other human V2-apex binding bnAbs. However, they do not have detectable tyrosine sulfation.

**Fig 7.**
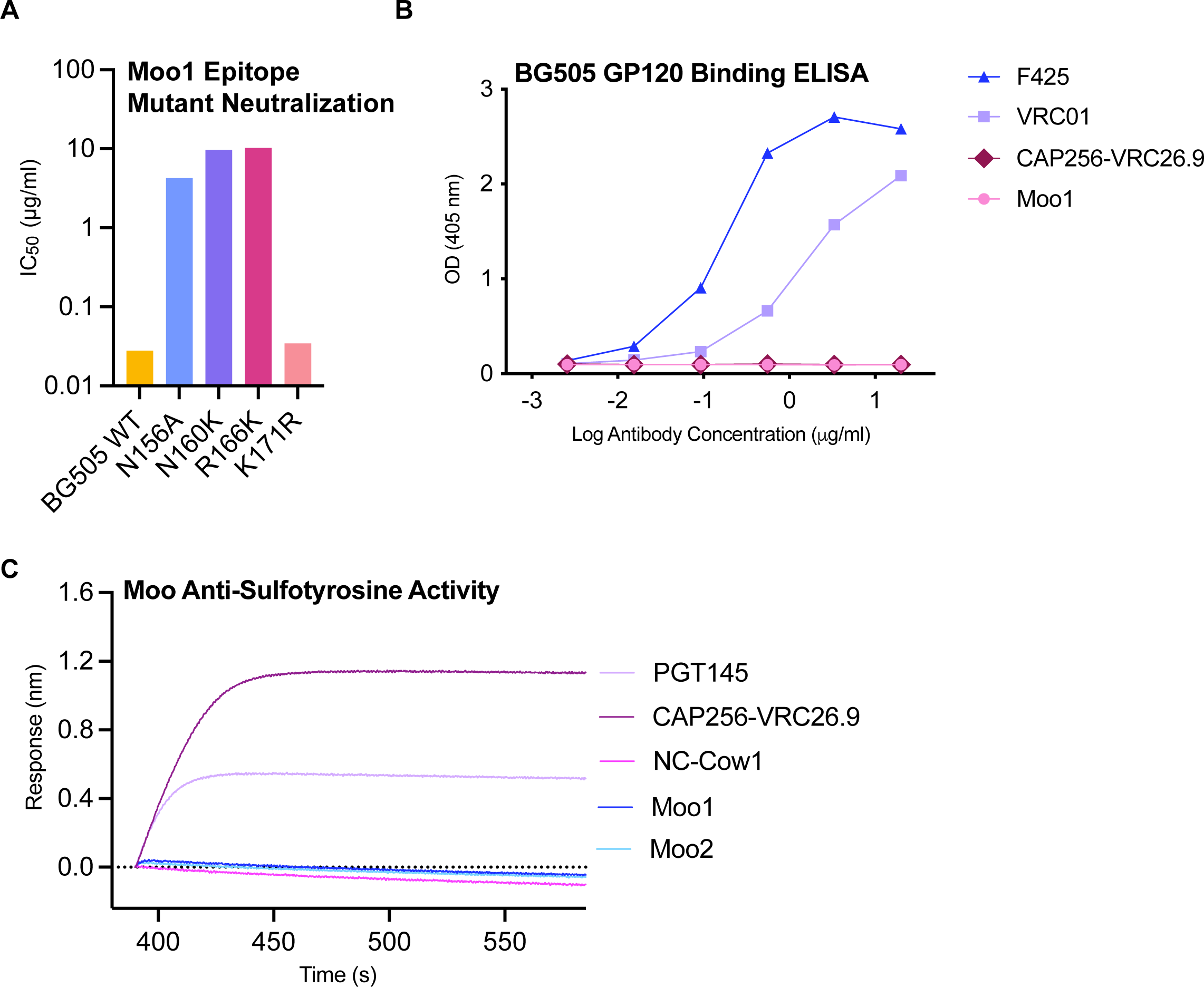
Broadly neutralizing antibodies Moo1 and Moo2 bind exclusively to trimer Env and not to monomeric gp120 and do not have detectable tyrosine sulfation. **(A)** Moo1 was tested for its ability to neutralize HIV BG505 V2-apex epitope-mutated virus. Bars show the measured IC50 (μg/ml) of the wild-type and mutated virus to show the decrease in neutralization potency at several key residues. **(B)** Moo1 was measured for binding to BG505 gp120 using ELISA. Positive antibody controls F425 and VRC01 are shown as well as negative control CAP256-VRC26.9. Moo1 and CAP256-VRC26.9 show no detectable binding to BG505 gp120. **(C)** Moo mAbs were evaluated for their ability to bind to a mouse derived anti-sulfotyrosine antibody using BLI. NC-Cow1 was used as negative controls. CAP256-VRC26.9 and PGT145 were used as positive controls.

## Discussion

Our previous studies demonstrated the ability of cows immunized with HIV well-ordered trimers to elicit broadly neutralizing serum antibody responses more rapidly than humans with natural infection or any wild-type model to date [27–29]. The cows immunized by Sok *et al.* 2017 demonstrated that a single immunogen, BG505, was able to elicit a broad and potent sera neutralizing response in a short amount of time [27]. After subsequent immunizations with the same trimer, monoclonal antibodies isolated from one of the cows were determined to be potent and broadly neutralizing antibodies and directed to the CD4bs [27]. Similarly, cows in Heydarchi *et al.* 2022 elicited a serum neutralizing response that was strengthened over time following immunization with multiple immunogens, including BG505 SOSIP, and again targeted the CD4bs [29]. One question not answered by the studies was whether immunization with monomeric BG505 gp120, which carries the native CD4bs and does bind cow CD4bs bnAbs, might be sufficient to induce cow CD4bs bnAbs. The answer to that question appears to be “no” in that, in contrast to Env trimer immunization, monomeric gp120 failed to show any evidence of bnAb induction. It appears likely that the hierarchy of favored epitopes in monomeric gp120 is very different from trimeric Env, and the induction of CD4bs Abs with the required features to access the CD4bs on native trimer is disfavored even in cows [39].

Our previous studies had also shown that, with the choice of appropriate trimer immunogens, V2-apex bnAbs could be induced in cows, although this took more immunizations, and the bnAbs were less potent and broad compared to those induced to the CD4bs by BG505 trimer [27, 28]. Here, we wished to see if, by mutation to greatly reduce bnAbs binding to the CD4bs, we could switch the induction of bnAbs to other sites using a mutated BG505 SOSIP trimer as an immunogen. Neutralization potency and breadth were reduced in the case of the mutated trimer and took longer to develop. The isolation of mAbs revealed two bnAbs directed to the Env V2-apex, which has many of the features of previously isolated cows and human V2-apex bnAbs [28, 38]. As previously found, these bnAbs made use of ultralong CDRH3s [27–29].

Finally, the potential for cow antibodies to inform mechanisms of recognition of the highly glycosylated HIV Env protein and to help design therapeutics continues to be an active area of study. The demonstrated stability and ability of cow ultralong bnAb NC-Cow 1 to bind gp120 in simulated vaginal fluid demonstrates the potential ultralong CDRH3 antibodies have as therapeutics, such as microbicides [27]. The isolated knobs and CDRH3 regions of ultralong antibodies have also been of interest for use as therapeutics outside of HIV in recent years.

They have been engineered for use as multivalent molecules, knobbody proteins, and functional knob-only proteins [40–46]. They have also been engineered to bind and potently neutralize SARS-CoV-2 as recombinant knob domains, highlighting their potential as anti-viral therapeutics [47].

## Acknowledgements

This work was supported by NIH grants GM105826 (VVS) and HD088400 (VVS) and the National Institute of Allergy and Infectious Diseases (NIAID) Consortium for HIV/AIDS Vaccine Development (CHAVD; UM1AI144462) (IAW, DRB, DS, and ABW).

## Author Contributions

DRB, DS, and VVS designed the immunization strategy. HS and WM performed cow immunizations, bleed, and processed sera and lymphocytes. PXA and MP sorting, sera neutralization assays, antibody isolation, screening, characterization, sera IgG purification. PXA was involved in strategy, analysis, and planning of experiments as well as BLI assays and competition assays. IAW provided SOSIP immunogens. GO, WHL, and ABW carried out planning, complexing, and negative stain imaging of monoclonal samples. PXA, DRB, and DS helped design and oversaw experiments. PXA, DRB, and DS wrote the manuscript.

## Materials and Methods

Cow Model: *Bos taurus*

### Ethics statement

The study was managed in accordance with the Public Health Service Policy on Humane Care and Use of Laboratory Animals as specified in the Health Research and Extension Act of 1985 (Public Law 99–158) or in accordance with the U.S Department of Agriculture policies as required by the Animal Welfare Act of 1966 (7.USC.2131 *et seq*) as amended in 1970, 1976, and 1985. This protocol (IACUC Protocol #3930.1) was approved by Texas A&M University Institutional Animal Care and Use Committee.

### Cow immunizations

*Bos Taurus* calves were primed and boosted to expand antigen specific B cell responses. Samples and calve groups were not randomized or blinded. The calves were bled from the jugular vein over the course of immunizations, and blood was collected to isolate PBMCs and sera from these time points. Immunizations and boosts were inoculated intradermally on both sides of the neck with a dose of 200 µg per calf (formulated in Montanide ISA 201 adjuvant: Seppic, France). Calves were immunized with BG505 SOSIP (Group 1) or BG505 D368R SOSIP (Group 2) on days 0, 36, 76, 118, 258, and 384. Calves immunized with BG505 gp120 (Group 3) were immunized on days 0, 36, 76, 118, and 258.

### Purification of sera for IgG

We heat-inactivated sera samples for 30 minutes using a 56°C water bath. Samples were then spun in a table-top centrifuge at max speed for 20 minutes. Sera samples were incubated, shaking overnight at 4°C with Protein G Sepharose beads (GE) and PBS in a 1:1:2 volume ratio. The following day, the beads were washed with 10x the volume of PBS. Beads were not allowed to dry out, and IgG was immediately eluted with IgG elution buffer (Pierce) and neutralized to a pH of 7.0 with 2M Tris pH 9.0. Eluted IgG was buffer exchanged into PBS and concentrated to the original sera volume, and filtered with a 0.45 µm filter.

### Pseudovirus production and neutralization Assays

HEK293T cells were co-transfected with plasmids of HIV Env and PSG3ΔEnv backbone in a 1:2 ratio for HIV pseudovirus production. DNA was combined with the transfection reagents FuGENE (Promega) and Transfectagro (Corning) or Opti-MEM. Cell cultures were incubated at 37°C for 72 hours before the supernatant was harvested, spun at 3000xg, and filtered with a 0.22 µM. Harvested pseudovirus was frozen at -80°C either neat or after concentrating with a 50k amicon concentrator and then frozen at -80°C. The frozen pseudovirus was titrated to determine the dilution needed for use.

Neutralization assays used 25 µl of pseudovirus and 25 µl of serially diluted monoclonal antibodies or IgG purified from sera for 1 hr at 37°C in a full-area 96-well tissue culture polystyrene microplate (Corning). After incubation, 20 µl of TZM-bl cells at a concentration of 0.5 million cells/mL with 40 µg/mL of dextran final [48]. Plates were incubated in a humidified 37°C incubator for 24 hr, and then 130 µl of media was added to all wells. 48-72 hr after cells were added, 120 µl of lysis buffer combined with Bright-Glo (Promega) was added to each well, and plates were read using a luminometer (BioTek). Cell controls and pseudovirus controls were done on each plate and averaged in each plate for analysis. Replicates were done on all samples and averaged. Purified IgG from sera samples were tested starting with a 1:35 dilution followed by 3-fold serial dilutions and reported as ID_50_ (1/dilution) titers. Monoclonal antibodies were diluted starting at 50 µg/ml and serially diluted 5-fold or 4-fold and reported as IC_50_ (µg/mL) titers. Analysis and graphing were done using Excel and Prism software.

### SOSIP trimer purification

SOSIP trimers and gp120 monomer were expressed in HEK293F cells and purified as described previously [49]. Briefly, SOSIP and gp120 were transfected with 150 µg of Furin DNA (SOSIP only), 300 µg of trimer DNA, and 1.5 mL of PEI MAX 40000 (Polysciences) or 1 mL of 293fectin in Transfectagro (Corning). Five days after transfection, supernatants were harvested by spinning at 3500xg and filtered with 0.22 or 0.45 µM filters. Supernatants were run over G*alanthus nivalis* lectin (Vector Labs) column or PGT145 affinity columns. As described previously, PGT145 affinity columns were made by PGT145 antibody coupled to CnBr-activated Sepharose 4B beads (GE) [49]. Purified proteins were run on size exclusion chromatography columns, specifically Superdex 200 10/300 GL column (Cytvia) in TBS or PBS. Some SOSIPs were biotinylated using Pierce Biotinylation Kit using the manufacturer’s protocol. Trimers were aliquoted into small volumes and frozen at -80°C. Antigenicity was tested for binding to known antibodies using ELISA or BLI.

### Single cell sorting

Cow PBMCs were sorted as previously described with study specific modifications [28, 50]. PBMCs were thawed until only a small ice piece remained in a 37°C bath and immediately added to 10mL of 37°C prewarmed 50% RPMI (Gibco) and 50% FBS buffer. Cells were then centrifuged at 400 xg for 5 min, and the supernatant was carefully discarded. 5mL of ice-chilled FACS Buffer (1% FBS, 1mM EDTA, 25 mM HEPES in 1x PBS) was added to the remaining cell pellet and gently resuspended. A small amount of cells were counted before again centrifuging and removing FACS Buffer and resuspended in 100 µL of FACS antibody master mix per 10 million cells for 30 min in the dark on ice. FACS antibody master mix was made with 3.75 µg of FITC labeled goat-anti cow antibodies (Abcam) and SOSIP baits diluted in FACS Buffer. 200 nM of each SOSIP bait was used per master mix. Biotinylated SOSIP baits were coupled to streptavidin-conjugated fluorophores and, when complexed, incubated with equal parts, 2x, or 3x competing Fabs, depending on trimer binding stoichiometry. Un-complexed BG505 was coupled with PE. Complexed BG505 was coupled with AF 647 and complexed with PGV04 Fab, PGT145 Fab, PGT151 Fab, or PGT121 Fab. All samples were separately stained and sorted by Fab complex. The second sort was done the same way but without antibodies complexed with SOSIP. BG505 was coupled with both BV421 and BUV737, and 25710 was coupled with PE. After 30 minutes, 1mL per 10 million cells of 1:300 diluted FVS510 Live/Dead stain (BD Biosciences) was added and incubated in the dark on ice for a further 15 min. Cells were resuspended in 10 mL of FACS buffer, centrifuged, and the pellet was resuspended again in 500 µl of FACS buffer per 10 million cells. The resuspended cell mixes were strained using a 5 mL round bottom FACS tube with a cell strainer cap (Falcon). Cells were sorted into dry 96-well plates and, once filled, immediately put on dry ice to freeze and transferred into a -80°C freezer. Data was analyzed using FlowJo.

### Single cell PCR amplification and cloning

Reverse transcriptase PCR (RT PCR) was used to generate cDNA. Each well received 0.225 µg of random hexamers (Gene Link), 0.33 mM dNTPs (New England Biolabs), 4.3 mM DTT, 15 µl of 3.33 mM RNaseOut (Invitrogen), 1x SuperScript IV Reverse Transcriptase Buffer (Invitrogen), 100 U of SuperScript IV Reverse Transcriptase Enzyme (Invitrogen), and RNAse Free Water until the final volume was met. RT PCR plates were run in thermocyclers using the following PCR program: 10min at 42°C, 10 min at 25°C, 10 min at 50°C, followed by 5 mins at 94°C. Generated PCR products were used in a nested PCR, PCR1, to amplify heavy chain and lambda light chain variable regions. Heavy chain PCR1 used primers

CowVHfwd1: CCCTCCTCTTTGTGCTSTCAGCCC,

CowIgGrev1: GTCACCATGCTGCTGAGAGA, and

CowIgGrev2: CTTTCGGGGCTGTGGTGGAGGC. PCR1s used 20 µl reactions per well/single cell using 3 µl of RT PCR product, 0.2 µM of forward primers total, and 0.2 µM of reverse primers total, 1x HotStarTaq Mastermix (New England Biolabs),the final volume was supplemented by RNAse Free Water. Thermocyclers were used with the following PCR program: 15s at 95°C, 49 cycles of 30s at 94°C, 30s at 55°C, and 60s at 72°C followed by an extension for 10 min at 72°C.

PCR2 to add tags for Gibson assembly used heavy chain primers

CowVHPCR2:

CATCCTTTTTCTAGTAGCAACTGCAACCGGTGTACATTCCMAGGTGCAGCTGCRGGAGTC and

CowIgGRevPCR2:

GGAAGACCGATGGGCCCTTGGTCGACGCTGAGGAGACGGTGACCAGGAGTCCTTGGCC.

Light chains were amplified with primers

L Leader 2 F: CACCATGGCCTGGTCCCCTCTG,

L Leader 15F: GGAACCTTTCCTGCAGCTC,

L Leader 16 F: GCTTGCTTATGGCTCAGGTC,

L Leader 35F: GACCCCAGACTCACCATCTC,

L Leader 45 F: AGGGCTGCGGGCTCAGAAGGCAGC,

L Leader 55 F: CTGCCCCTCCTCACTCTCTGC, and

Cow_LC_rev1: AAGTCGCTGATGAGACACACC. PCR2 was done for the light chain using primers

Fwd-PCR2-LC: CATCCTTTTTCTAGTAGCAACTGCAACCGGTGTACACCAGGCTGTGCTGACTCAG and LC_REV_const-PCR2: GTTGGCTTGAAGCTCCTCACTCGAGGGYGGGAACAGAGTG. PCR Products from PCR 2 were used for 25 µl reactions per well/single cell using 2 µl of PCR product from PCR1, 0.2 mM dNTPs at 10 mM each (New England Biolabs), 1 µM of forward primers total, 1 µM of reverse primers total,1.5 mM MgCl_2_, 1x HF Buffer (Thermo Fisher), 0.5 U of Phusion Enzyme (Thermo Fisher), and supplemented with RNAse Free Water until the final volume was achieved. PCR 2 used the following PCR program: 30s at 98°C then 34 cycles of 10s at 98°C, 40s at 72°C, followed by an extension for 5 min at 72°C.

To determine heavy chain or light chain amplification, PCR products were resolved using E-gel 96 Agarose Gels 2% (Invitrogen). SPRIselect beads (Beckman Coulter) were used to purify successful amplifications. NEBuilder HiFi DNA assembly system was used to assemble cow variable regions with human antibody expression vectors and appropriate IgG1 or Ig lambda constant domains. 10 ng of restriction enzyme digested backbone and 20 ng of SPRI purified PCR variable regions were combined with an appropriate volume of 2x HiFi master mix (New England Biolabs) for one hour at 50°C. Assembled products were transformed into DH5 Alpha Competent Cells (Biopioneer) and grown overnight at 37 °C. Single colonies were picked and purified using the QIAprep Spin Miniprep Kit (Qiagen). Products were sanger sequences and analyzed using International ImMunoGeneTics (IMGT) Information System (www.imgt.org) V-quest [51].

### Large-scale antibody production and purification

Monoclonal antibodies were transiently expressed in 293F cells. Heavy and light chain DNA was added in a 1:1 ratio with 1.5 mL PEI: 1mL cell volume as a transfection reagent and diluted in Opti-MEM (Thermo Fisher) or Transfectagro (Corning) media. Five days after transfection, supernatants were harvested and sterile filtered through a 0.22 or 0.45 µm filter. IgG containing supernatant was batch bound overnight at 4°C using Praesto AP (Purolite). The next day, beads were washed 10x with PBS, and the antibody was eluted using 0.2 M citric acid and neutralized to pH 7 with 2M Tris pH 9.0. Antibodies were buffer exchanged into PBS.

Fabs were papain digested using Pierce Fab Preparation Kit (Thermo Fisher) following the manufacturer’s protocol for human Fab digest. Polyclonal IgG from sera were also prepared using Pierce Fab Preparation Kit with each digest at 0.5 mg/mL and a digest time of 10-16 hr.

### High throughput antibody screening

High-throughput screening of antibodies was done using Expi293 cells grown in 3 mL plates in 24 deep-well plates (Thermo Fisher). Cultures were grown for four days, harvested, and neat ELISA assays were used to test for expression and SOSIP binding. Antibodies that expressed were further screened for neutralization as previously described using neat supernatant. If neutralization was detected, antibodies were expressed in 30 to 50 mL cultures and purified using Praesto AP (Purolite) beads as previously described.

### BioLayer interferometry (BLI) assays

BLI competition assays were performed using an Octet RED384 instrument at 30°C in a 96-well polypropylene black 96-well microplate (Greiner). Antibody competition assays were done using an in-tandem assay. 222 nM of 6xHis-tagged BG505 SOSIP was immobilized with nickel-charges tris-NTA (Ni-NTA) biosensors (Sartorius) for 10 min and transferred to PBS for 1 min to wash off unbound SOSIP. Sensors were then moved into monoclonal Moo1 and Moo2 antibodies at a concentration of 100 µg/ml for 10 min and washed off in PBS for 1 min. Biosensors were then moved into known antibodies at a concentration of 25 µg/ml for 5 min. The percent inhibition in binding was calculated with the formula: Percent binding inhibition (%) = 1-(competitor antibody binding response in the presence of saturating antibody) / (binding response of the competitor antibody without saturating antibody). BLI assays to detect tyrosine sulfation by binding antibodies with anti-sulfotyrosine antibodies were performed using polypropylene black 384-well microplate (Greiner) at 30°C in octet buffer (0.05% Tween in PBS). Monoclonal Moo1 and Moo2 antibodies were immobilized on anti-human IgG Fc Capture (AHC) biosensors (Sartorius) to capture monoclonal antibodies at a concentration of 50 µg/mL for 5 min and washed with octet buffer. Biosensors were then moved to 1:50 diluted mouse Anti-Sulfotyrosine Antibody, Clone Sulfo-1C-A2 (MilliporeSigma), to detect binding for 200 sec. All dilution and incubation steps were performed in 1x PBS with 0.05% Tween. Analysis and graphing were done using Prism software.

### ELISA assays

All ELISA assays were done with half-area 96-well high binding ELISA plates in 50 µl per well, except blocking which was done in 150 µl/well (Corning). Washes were done three times using PBS containing 0.05% Tween20, and dilutions were done in PBS containing 1% BSA and 0.025% Tween20 unless otherwise stated. All plates were developed with phosphatase substrate (Sigma-Aldrich) diluted in alkaline phosphatase staining buffer (pH 9.8), according to the manufacturer’s instructions, and read with 405 nm optical density using a microplate reader (Molecular Devices). All data and graphs were analyzed using Prism software.

Sera competition ELISA assay plates were coated with PBS containing 250 ng of anti-6x His tag monoclonal antibody (Invitrogen) and left covered overnight at 4°C. Plates were then washed and blocked with 3% BSA for 1 hour at room temperature. After washing plates, 125 µg of his-tagged BG505 was diluted and added to all wells for 1 hr. IgG from sera was serially diluted and incubated in the plate for 1 hr and washed. Biotinylated monoclonal anti-HIV antibodies were added at their constant EC_70_ value for 1 hr and washed. Secondary alkaline phosphatase conjugated to streptavidin at a dilution of 1:1000 (Jackson ImmunoResearch) was added to plates and allowed to incubate for 1 hr then washed. A substrate was added, and plates were allowed to develop and then be read. Biotinylated antibody EC_70_ values were measured using the same methods but without the step with purified IgG from sera and serially diluted.

ELISA assays for gp120 binding used plates coated with PBS containing 250 ng of BG505 gp120 and left covered overnight at 4°C. The following day, plates were washed and blocked with 3% BSA for 1-2 hrs. Monoclonal antibodies were added using serial dilutions starting at 20 µg/mL and incubated for 1 hr. After washing, alkaline phosphatase conjugated anti-human F(ab’)_2_ secondary antibodies (Jackson ImmunoResearch) diluted 1:1000 and added to plates to incubate for 1 hr and then washed. Substrate was then added, and plates were read as mentioned previously.

High throughput screened antibodies were evaluated for antibody expression with an ELISA. Plates were coated overnight at 4°C with 100 ng of AffiniPure Fragment Goat Anti-Human IgG, F(ab’)₂ (Jackson ImmunoResearch). The plates were washed the next day and blocked with 3% BSA for 2 hr at 37°C. Neat supernatant was added to the plate for 1 hr. After a wash step, the secondary antibody, 1:1000 diluted Alkaline phosphatase conjugated anti-human Fc antibodies (Jackson ImmunoResearch), was incubated in the plate for 1 hr and then washed. Substrate was then added, and plates were read as mentioned previously.

High throughput screened antibodies were evaluated for BG505 SOSIP binding with an ELISA. Plates were coated overnight at 4°C with 100 ng of 6x His tag monoclonal antibodies (Invitrogen) in PBS and washed the next day. Plates were blocked with 3% BSA for 2 hr at 37°C and then washed. Each well received 125 µg of 6x His-tagged BG505 SOSIP and was incubated for 1 hr and then washed. Neat supernatant was added to the plate for 1 hr and then washed. After a wash step, the secondary antibody, 1:1000 diluted Alkaline phosphatase conjugated anti-human F(ab’)_2_ antibodies (Jackson ImmunoResearch) were incubated in the plate for 1 hr and then washed. Substrate was then added, and plates were read as mentioned previously.

### Negative-stain electron microscopy

Trimer was complexed with Moo1 Fab and RM20A3 base Fab at 3x molar excess Fab:trimer overnight at room temperature. The protein complex was then diluted to 0.02 mg/ml in 1x Tris-buffered saline, and 3 µL were applied to a 400 mesh Cu grid, blotted with filter paper and stained with 2% (w/v) uranyl formate. Micrographs were collected on a 120 keV FEI Tecnai TF20 microscope with a TVIPS TemCam F416 CMOS camera (1.68 Å/pixel; 62,000x magnification) using Leginon[52]. Particles were picked using DoGPicker and 2D and 3D classifications and 3D refinement was performed using Relion 3.0 [53, 54]. The final map was segmented in UCSF Chimera to generate figures, and deposited to the Electron Microscopy Data Bank (EMDB) [55].

